# Partner-specific induction of *Spodoptera frugiperda* immune genes in response to the entomopathogenic nematobacterial complex *Steinernema carpocapsae-Xenorhabdus nematophila*

**DOI:** 10.1101/800656

**Authors:** Louise Huot, Audrey Bigourdan, Sylvie Pagès, Jean-Claude Ogier, Pierre-Alain Girard, Nicolas Nègre, Bernard Duvic

## Abstract

The *Steinernema carpocapsae-Xenorhabdus nematophila* association is a nematobacterial complex (NBC) used in biological control of insect crop pests. The ability of this dual pathogen to infest and kill an insect strongly depends on the dialogue between the host’s immune system and each partner of the complex. Even though this dialogue has been extensively studied from the two partners’ points of view in several insect models, still little is known about the structure and the molecular aspects of the insects’ immune response to the dual infection. Here, we used the lepidopteran pest *Spodoptera frugiperda* as a model to analyze the respective impact of each NBC partner in the spatiotemporal immune responses that are induced after infestation. To this aim, we first analyzed the expression variations of the insect’s immune genes in the fat bodies and hemocytes of infested larvae by using previously obtained RNAseq data. We then selected representative immune genes for RT-qPCR investigations of the temporal variations of their expressions after infestation and of their induction levels after independent injections of each partner. We found that the fat body and the hemocytes both produce potent and stable immune responses to the infestation by the NBC, which correspond to combinations of bacterium- and nematode-induced ones. Consistent with the nature of each pathogen, we showed that *X. nematophila* mainly induces genes classically involved in antibacterial responses, whereas *S. carpocapsae* is responsible for the induction of lectins and of genes expected to be involved in melanization and cellular encapsulation. In addition, we found that two clusters of unknown genes dramatically induced by the NBC also present partner-specific induction profiles, which paves the way for their functional characterization. Finally, we discuss putative relationships between the variations of the expression of some immune genes and the NBC’s immunosuppressive strategies.

**Author summary:** Entomopathogenic nematodes (EPNs) are living in the soil and prey upon insect larvae. They enter the insect by the natural orifices, and reach the hemocoel through the intestinal epithelium. There, they release their symbiotic bacteria that will develop within the insect and eventually kill it. Nematodes can then feed and reproduce on the insect cadaver. By using transcriptomic approaches, we previously showed that Lepidoptera larvae (caterpillars of the fall armyworm *Spodoptera frugiperda*) produce a strong immune response in reaction to infestation by EPNs. However, we do not know if this immune reaction is triggered by the nematode itself -*Steinernema carpacapsae* - or its symbiotic bacteria - *Xenorhabdus nematophila*. To answer this question, we present in this work a careful annotation of immunity genes in *S. frugiperda* and surveyed their activation by quantitative PCR in reaction to an injection of the bacteria alone, the axenic nematode or the associated complex. We found that the immune genes are selectively activated by either the bacteria or the nematode and we discuss the implication of which pathway are involved in the defense against various pathogens. We also show that a cluster of newly discovered genes, present only in Lepidoptera, is activated by the nematode only and could represent nematicide genes.

## Introduction

The *Steinernema-Xenorhabdus* nematobacterial complexes (NBCs) are natural symbiotic associations between nematodes and enterobacteria that are pathogenic for insects. The soil-living nematodes infest insects through the respiratory and/or the intestinal tract (1) and reach the hemocoel, the internal body cavity, where they release their intestinal symbionts. The bacteria then grow extracellularly in the hemolymph, the insect equivalent of blood, and improve the nematodes’ pathogenicity as well as their ability to reproduce in the host dead body (2). Until now, about 90 species of *Steinernema* have been identified, among which several are usable as biological control agents against diverse insect crop pests (3, 4). In consequence, their interactions with insects have been extensively studied for about 50 years (5). These studies have shown that in addition to ecological and morphological parameters (3), the NBCs’ interactions with the host’s immune system is one of the most crucial factors influencing their ability to infest and kill a given insect (6–8).

Insects possess an elaborate immune system, which is able to respond by adapted ways to diverse types of pathogens and of infections. This system firstly relies on protective external barriers such as the cuticle, or the peritrophic matrix in the midgut (9, 10). It then relies on local defenses of the surface epitheliums, which repair efficiently (11–13) and produce toxic factors such as antimicrobial peptides (AMPs) (14–17) and reactive oxygen species (18). The third line of defense of insects is provided by the hemocytes, which are the circulating immune cells. They can produce diverse types of immune responses, including AMP synthesis, phagocytosis, nodulation, encapsulation, coagulation and melanization (19). Nodulation and encapsulation are cellular immune responses respectively consisting in the engulfment of bacterial aggregates and of large invaders via hemocytes aggregation (19). Together with coagulation, these responses are coupled with a melanization process consisting in series of phenolic compounds oxidations resulting in synthesis of reactive molecules and melanin that participate of pathogens trapping and killing (20, 21). Finally, the fat body, a functional equivalent of the mammalian liver, produces potent systemic humoral immune responses involving a massive secretion of AMP cocktails in the hemolymph. These responses can be induced by two major signaling pathways of insect immunity; the Imd pathway, which is mainly activated by Gram negative bacteria, and/or the Toll pathway, which is mainly activated by Gram positive bacteria, fungal organisms and by proteases released by pathogens (22, 23).

The *Steinernema-Xenorhabdus* NBC whose interactions with the immune system have been the most extensively studied is the *S. carpocapsae-X. nematophila* association. These interactions have firstly been studied from the NBC point of view, which allowed the identification of a multitude of immunoevasive and immunosuppressive strategies. For instance, studies in *Rhynchophorus ferrugineus* and *Galleria mellonella* have respectively shown that the cuticle of *S. carpocapsae* is not recognized by the host’s immune system (24, 25) and that the nematode secretes protease inhibitors impairing the coagulation responses (26, 27). Studies in diverse insect models have also shown that both partners produce factors impairing melanization (28–31), hemocyte’s viability (32–36) and the production of cellular immune responses by several ways (27–29, 31, 37, 38). Finally, both *X. nematophila* and *S. carpocapsae* secrete proteolytic factors degrading cecropin AMPs (39, 40) and the bacterium has also been shown to reduce more globally the hemolymph antimicrobial activity, as well as AMP transcription in lepidopteran models (24, 39, 41, 42).

On the other hand, the description of these interactions from the hosts’ points of view is at its beginning. This aspect has mainly been studied in the *Drosophila melanogaster* model, with a first transcriptomic analysis of the whole larva responses to infestations by entire NBCs and by axenic nematodes (43). This analysis has shown that several immune processes are induced by both pathogens at the transcriptional level. For instance, the authors found in each case an overexpression of genes related to the Imd and Toll pathways that was accompanied by the induction of a few AMP genes. They also found an upregulation of genes related to melanization, coagulation, or involved in the regulation of cellular immune responses (43). Complementary gene knockout experiments in this model demonstrated an involvement of the Imd pathway in the response against *X. nematophila* (44) and revealed a possible involvement of the Imaginal Disc Growth Factor-2, the intestinal serine protease Jonah 66Ci (45) as well as TGF-β and JNK pathways members in the regulation of anti-nematode immunity (46, 47).

In order to improve our understanding of the dialogue that takes place between this NBC and its host, we recently published a topologic transcriptomic analysis of the response of the lepidopteran model *Spodoptera frugiperda* to the infestation (48). This analysis was focused on the three main immunocompetent tissues that are confronted to the NBC, which are the midgut (the main entry site in the hemocoel), the hemocytes and the fat body. The RNAseq experiment showed that there was no potent or well-defined transcriptional response in the midgut. However, we observed dramatic transcriptional responses in the fat body and the hemocytes at 15 h post-infestation, which is a middle time point of the infection. In agreement with the results obtained in *D. melanogaster* whole larvae (43), global analysis of these responses showed they are dominated by immune processes. The objective of the present study is to go further in the analysis of these induced immune responses. In order to describe them with high accuracy, we first examine the expression variations of all the immune genes that have been identified in the insect’s genome. We then use tissue RT-qPCR experiments to analyze the temporal dynamics and the relative contribution of each NBC partner in the identified immune responses. Our results show that a large number of immune genes are responsive in either one or the two tissues during the infestation, with activation of antimicrobial and cellular immunities, of melanization, coagulation and of metalloprotease inhibition. These responses were found to be stable over the time post-infestation and to consist in combinations of *X. nematophila*-induced and *S. carpocapsae*-induced responses in each tissue. The *X. nematophila*-induced responses mainly correspond to genes that are classically involved in antibacterial immunity, whereas the *S. carpocapsae*-induced ones mainly include lectins and genes potentially involved in melanization and encapsulation. In addition, our RT-qPCR experiments show that two previously identified candidate clusters of uncharacterized genes (48) also present partner-specific induction profiles. Our hypothesis is that they may correspond to new types of anti-nematode and antibacterial immune factors found in *Spodoptera* genus and lepidopteran species, respectively.

## Results & Discussion

### Hemocytes’ and fat body’s immune responses

In order to get an accurate picture of the *S. frugiperda* transcriptional immune responses to the NBC infestation, we first used a previously published list of immune genes identified by sequence homology in the *S. frugiperda* genome (49). We then looked at their expression variations in the fat body and in the hemocytes (S1A Table) and we completed the repertoire with additional putative immune genes that we directly identified from our RNAseq data (S1B Table). In total, we present the annotation of 226 immune or putative immune genes of which 132 were significantly modulated at 15 h post-infestation (hpi) (Sleuth, p-value < 0.01; |Beta| > 1; all count values > 5 in at least one condition) in one or both tissues (Fig 1). Among them, 62 were involved in antimicrobial responses (Fig 1A), 18 were related to melanization (Fig 1B), 23 were involved in cellular responses (Fig 1C) and the 29 remaining genes were grouped in a category called “diverse” due to pleiotropic or poorly characterized functions (Fig 1D).

**Figure 1.**
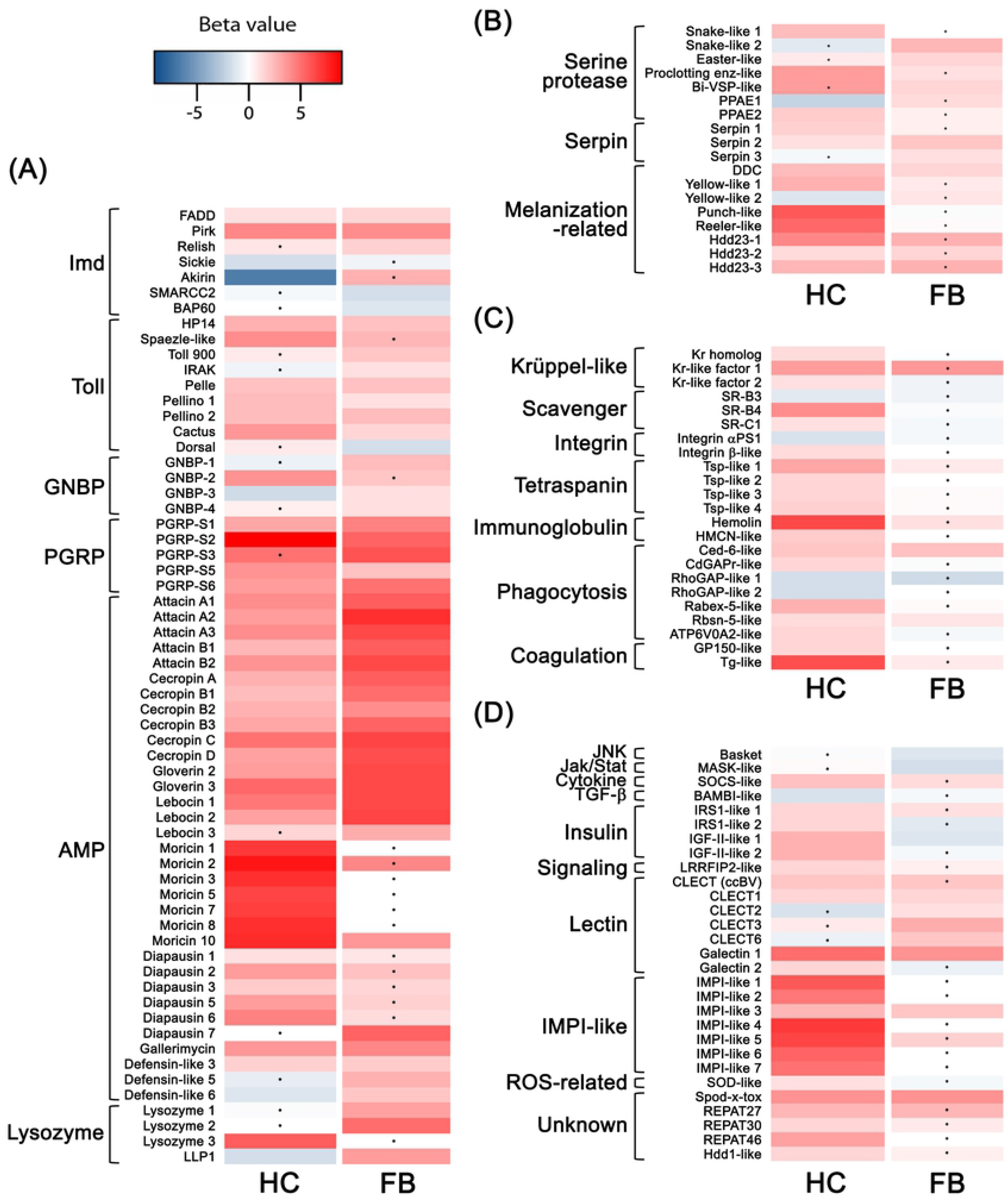
Expression variations of the differentially expressed immune genes after infestation by the nematobacterial complex. Heatmaps showing the expression variations of the differentially expressed immune genes in the hemocytes and in the fat body at a middle time point of 15 h post-infestation. Infestation experiments were performed in triplicate with N=9 larvae per sample. RNAseq data were analyzed with the Kallisto/Sleuth softwares using statistical thresholds of 0.01 for p-values, −1 and +1 for Beta value (biased equivalent of log2 fold change) and 5 for pseudocount means. The immune genes were identified by homology and classified as (A) antimicrobial immunity-related, (B) melanization-related, (C) cellular immunity-related and (D) diverse immune responses. Black dots indicate genes with statistically non-significant variations to the controls in the corresponding tissue; HC : Hemocytes, FB : Fat body.

#### Antimicrobial responses

In the antimicrobial response category, 58 genes were found to be upregulated in at least one of the two tissues (Fig 1A). The signaling genes encoded 3 and 8 members of the Imd and Toll pathways, respectively, as well as 5 short catalytic peptidoglycan recognition proteins (PGRP-S), which are probably involved in the regulation of these pathways by peptidoglycan degradation (50, 51) (Fig 1A). Four other genes were considered as involved in recognition. They encoded Gram negative binding proteins (GNBPs), which have been reported to recognize peptidoglycans or β-glucans and participate in the further activation of the Toll pathway (22) (Fig 1A). Finally, the effector genes encoded 33 antimicrobial peptides (AMPs) belonging to all the *S. frugiperda*’s AMP families (49) plus 4 lysozymes and lysozyme-like proteins (LLPs) (Fig 1A). Depending on their families and on the insect species, AMPs can present varied activity spectra, ranging from antiviral or antibacterial activities to anti-fungal and anti-parasitic ones (52). Varied activity spectra have also been found for several insects’ lysozymes and LLPs (53–57). Interestingly, all of the categories and subcategories cited above were represented in the two tissues, indicating that their antimicrobial responses are diversified and that the factors responsible for their disappearance in the hemolymph (24, 41) probably act at a post-transcriptional level. About a half of the genes presented similar and significant induction profiles in the hemocytes and in the fat body. This is for instance the case of the usually anti-Gram negative bacteria attacin, cecropin and gloverin AMPs (52), which were all highly induced in the two tissues (Fig 1A), suggesting they both respond to the bacterial partner *X. nematophila*. On the other hand, all the induced GNBP, lysozyme and LLP genes were found to be either significantly induced in the hemocytes or in the fat body, and in the AMP category, tissue-specificities were observed for diapausin, defensin-like and most moricin genes (Fig 1A).

Only 8 antimicrobial response genes were found to be significantly downregulated (Fig 1A). Interestingly, 4 were involved in the Imd pathway whereas the 4 remaining ones were dispersed between the AMP, GNBP and lysozyme categories (Fig 1A). The Imd pathway downregulated factors included *sickie* and the *akirin* in the hemocytes and *SMARCC2* and *BAP60* in the fat body (Fig 1A). In *D. melanogaster*, Sickie participates in the activation of Relish, the transcription factor of the Imd pathway (58) and the akirin acts together with the Brahma chromatin-remodeling complex, containing BAP60 and SMARCC2, as cofactor of Relish to induce the expression of AMP genes (59). Given the potent induction of anti-Gram negative bacteria immune responses in the two tissues, the down-regulation of these genes could be attributed to immune regulations. However, it has been shown that in the close species *S. exigua*, live *X. nematophila* reduces the expression of several AMP genes, including attacin, cecropin and gloverin (42, 60, 61). It would thus be of particular interest to determine whether the observed down-regulations are related to this immunosuppressive effect.

To summarize, the antimicrobial responses are potent and diversified in the two tissues, with a common induction of genes that probably respond to *X. nematophila*. Yet unexplained tissue-specific responses were observed and the results show a down-regulation of Imd pathway members that could be related to a previously described transcriptional immunosuppressive effect of the NBC. However, this effect would not be potent enough to suppress the humoral responses at this time point, suggesting that the NBC probably uses other immunosuppressive strategies in this model.

#### Melanization

In the melanization category, 16 genes were found to be upregulated in at least one of the two tissues (Fig 1B). These genes firstly encoded 6 serine proteases (Fig 1B) that were considered as members of the prophenoloxidase (proPO) system. The proPO system is an extracellular proteolytic cascade ending in the maturation of the proPO zymogen into PO, which initiates the melanization process (62). Among the upregulated serine proteases, PPAE2 is the only one that is known to take part in proPO processing whereas the other proteases were included in this category because of their characteristic CLIP domains and of their low homology with the serine proteases acting upstream of the Toll pathway in *D. melanogaster* (63). The other upregulated genes in this category included 3 serpins, which are known to regulate the proPO system in several model insects (62), 3 melanization enzymes, DDC, Yellow-like 1 and Punch-like (64, 65) as well as 4 genes, Reeler-1 and 3 Hdd23 homologs, that are involved in melanization and nodule formation in other models (66, 67) (Fig 1B). Despite of tissue-specific induction patterns, serine proteases and serpins were found in the two tissues (Fig 1B), suggesting that both participate in the stimulation of the proPO system, which is consistent with results obtained in other interaction models (68–70). However, with the exception of the DDC, all the melanization enzymes as well as the nodulation-related genes were specifically induced in the hemocytes (Fig 1B), which is consistent with the very localized nature of this immune response (65) that is mainly mediated by hemocyte subtypes.

Finally, only 2 genes, PPAE1 and Yellow-like 2, were found to be significantly down-regulated in this category (Fig 1B). Both were specifically repressed in the hemocytes, which could be due to functional interferences with their upregulated homologs (PPAE2 and Yellow-like 1).

In summary, our results suggest that both the hemocytes and the fat body participate in induction and regulation of melanization in response to the NBC and no sign of transcriptional immunosuppression is detected for this response. These results are in agreement with the previous identification of diverse PO inhibitors in both *S. carpocapsae* (28, 29) and *X. nematophila* (30, 31).

#### Cellular responses

In the hemocytes, 19 upregulated genes were placed in the cellular responses category (Fig 1C). The signaling ones encoded 3 homologs of the transcription factor Krüppel (Kr) (Fig 1C). In *D. melanogaster*, Kr and Kr homologs are involved in several developmental processes such as embryo patterning (71), organogenesis (72–74), and cell differentiation (75). More specifically in the hemocytes, Kr has been shown to take part in hemocytes’ differentiation and/or activation (76), a crucial step for the induction of cellular immune responses. The recognition genes encoded 3 cellular receptors of the Scavenger (SR) and Integrin families plus the hemolin, a secreted immunoglobulin-containing protein (Fig 1C). Both Scavenger receptors and integrins are known to act as membrane receptors in phagocytosis of bacteria and apoptotic cells (77). In addition, integrins are involved in diverse processes, including cell motility and adhesion, and encapsulation (78, 79). The hemolin is known to act as an opsonin by increasing phagocytosis and nodulation of bacteria in *Manduca sexta* (80). Among the effector genes, we first identified 5 upregulated genes corresponding to conserved intracellular phagocytosis-related proteins. They included Ced-6, the Rabenosyn-5 (Rbsn-5-like), a V-ATPase subunit (ATP6V0A2-like) and 2 small GTPase Activating Proteins (Rabex-5-like, CdGAPr-like) (77) (Fig 1C). We also found genes encoding membrane proteins, such as the immunoglobulin-containing hemicentin (HMCN-like) (81) and 4 tetraspanin-like (Tsp-like) proteins (82) (Fig 1C), that could participate in cell-cell adhesion and cellular immune responses. Interestingly, one of the upregulated tetraspanins (Tsp-like 3) presented 79.5% identity with the *Manduca sexta* (Lepidoptera : Noctuidae) tetraspanin D76, which takes part in hemocytes aggregation during capsule formation by trans-interacting with a specific integrin (83). Finally, 2 genes encoding proteins similar to the *D. melanogaster* clotting factors GP150 (84) and a transglutaminase (Tg-like) (85) were also found upregulated (Fig 1C). Only 2 genes (Ced-6-like, Rbsn-5-like) of the cellular responses category were found to be upregulated in the fat body (Fig 1C) and both encoded intracellular proteins that are probably not related to immunity in this tissue.

All the 4 down-regulated putative cellular immunity-related genes were specifically modulated in the hemocytes (Fig 1C). They encoded 2 Rho GTPase Activating Proteins (RhoGAP-like), a scavenger receptor similar to the *D. melanogaster* Croquemort receptor (SR-B3) and a homolog of the *D. melanogaster* integrin α-PS1. In *D. melanogaster*, Croquemort has been shown to take part in phagocytosis of apoptotic cells and of the Gram positive bacterium *Staphylococcus aureus* but not of the Gram negative bacterium *Escherichia coli* (79, 86). Integrin α-PS1 is a ligand of the extracellular matrix protein laminin (87). It is involved in migration and differentiation of several cell types during development (88–90) but does not seem to be required for any immune process. Their down-regulations are thus probably due to their uselessness in the context of the response to the NBC.

Overall, the results suggest that all types of cellular responses are transcriptionally induced at 15 hpi, including phagocytosis and nodulation, as well as encapsulation that would be adapted to the bacterial partner or the nematode, respectively. In addition, the induction of coagulation responses is particularly interesting, since many clotting factors participate in *D. melanogaster* resistance to infestation by another type of NBC, the *Heterorhabiditis bacteriophora-Photorhabdus luminescens* association (91–94). Moreover, despite *S. carpocapsae* does not pierce the insects’ cuticles as *H. bacteriophora* (1), it has been shown to express at least two secreted proteases with inhibitory activities towards the formation of clot fibers and coagulation-associated pathogen trapping (26, 27). Once again, the induction of such immune responses is consistent with the previous identification of several virulence factors of the NBC targeting cellular immunity (26, 28, 29, 31–38).

#### Diverse immunity-related genes

A total of 29 modulated genes were involved in other diverse immune processes. They included 10 up- or down-regulated signaling genes, 7 upregulated recognition genes, 8 upregulated effector genes and 5 upregulated genes of unknown functions that are known to be modulated after immune challenge (Fig 1D).

The signaling genes firstly encoded 2 insulin-like growth factor (IGF-II-like) and 2 insulin receptor substrate homologs (IRS1-like) (Fig 1D). Insulin signaling is known to have a deleterious impact on the induction of systemic immune responses in the fat body of *D. melanogaster* (95) whereas insulin increases hemocyte proliferation in the hemolymph of mosquitoes (96) as well as in the hematopoietic organs of the lepidopteran model *Bombyx mori* (97). In agreement with these assertions, we found that 2 of these genes were down-regulated in the fat body, but all 4 genes were upregulated in the hemocytes (Fig 1D). Two other signaling genes were found to be specifically overexpressed in the hemocytes. The first one is a homolog of the *Litopenaeus vannamei* (Decapoda: Penaeidae) leucine-rich repeat flightless-I-interacting protein 2 (LRRFIP2-like) (Fig 1D), which has been shown to upregulate AMP expression in *L. vannemei* as well as in *D. melanogaster* (98). On the other hand, 3 signaling genes were found to be strictly down-regulated (Fig 1D). Interestingly, these genes included a member of the TGF-β pathway (BAMBI-like) in the hemocytes and a member of the JNK pathway in the fat body (Basket), two pleiotropic pathways that are currently suspected to take a part in the *D. melanogaster* immune response to nematodes after NBC infestation (47, 99–101). The third down-regulated gene was found in the fat body and encoded MASK, an inducer of the Jak/Stat pathway (102). In the fat body, the Jak-Stat pathway has mainly been shown to induce the expression of cytokines (103) and of a putative opsonin belonging to the TEP family (104). Remarkably, several Tep genes have been shown to participate in antibacterial immunity after NBC infestation in *D. melanogaster* (91, 105–107). All of these down-regulations could thus impair the insect’s immune response to the NBC. However, more detailed analyses of their functions and modulations would be required to hypothesize immunosuppressive effects of the NBCs.

All 7 upregulated recognition genes encoded lectins (Fig 1D). Five of them encoded C-type lectins (CLECT), which are known to be involved in binding of diverse pathogens (108), including bacteria and nematodes (109). This binding can then stimulate several immune responses, such as bacterial aggregation, melanization, phagocytosis, nodulation and encapsulation (108). The 2 others encoded galectins, which are involved in diverse aspects of mammalian immunity, including pathogens binding (110), and are considered as relevant candidate immune proteins in insects (111). Despite a larger set of upregulated lectins was identified in the fat body, members of these protein families were found upregulated in the two tissues.

In the hemocytes, the upregulated effector genes firstly encoded a homolog of the superoxide dismutase (SOD-like), a conserved detoxifying enzyme involved in responses to reactive oxygen species (112) (Fig 1D). The 7 remaining genes encoded proteins with similarity to insect metalloproteinase inhibitors (IMPI-like) (Fig 1D), whose functions have only been studied in the lepidopteran model *Galleria mellonella*. The only characterized IMPI encodes two proteins of which one is probably involved in the regulation of extracellular matrix remodeling and the second specifically targets metalloproteinases from pathogens (113, 114). *S. carpocasape* and *X. nematophila* both express several secreted serine proteases as well as metalloproteinases during the infectious process (39, 115–120). The induction of such immune responses could interfere with some of these proteinases to impair the NBC’s virulence and/or survival. Interestingly, all but one of these IMPI homologs were found to be specifically upregulated in the hemocytes, a tissue-specificity that had not been highlighted in previous reports (121, 122).

Finally, the remaining genes of unknown function encoded Spod-x-tox, a protein without antimicrobial activity which contains tandem repeats of defensin-like motifs (123), 3 REPAT genes, which are known to be induced in the midgut after exposure to toxins, viruses and intestinal microbiota perturbations in the close species *S. exigua* (124–126), and Hdd1, which is induced in response to bacteria and peptidoglycan in the lepidopteran models *Hyphantria cunea* and *Bombyx mori* (127, 128) (Fig 1D).

In summary, we found an important additional mobilization of several relevant candidate immune genes, including mainly insulin signaling factors and IMPIs in the hemocytes and lectins in the fat body. In addition, these results suggest that the candidate immune pathways TGF-β, JNK and Jak/Stat could be down-regulated. Such down-regulations are in disagreement with the results of Yadav and colleagues (43) in *D. melanogaster* and thus would require further investigation.

### Temporal analysis of the induced immune responses

In order to put the *S. frugiperda* immune responses in relation with the infectious process, we then described their temporal dynamics in each analyzed immunocompetent tissue. To this aim, we monitored with RT-qPCR experiments the induction levels of selected representative immune genes from 5 hpi, the mean time at which nematodes release *X. nematophila* in the hemocoel, to 20 hpi, which is about 9 hours before the first insect deaths (S1 Fig).

In the hemocytes, the selected genes included 15 genes of the antimicrobial response, 2 genes involved in melanization, 5 cellular response genes, 2 lectins and one IMPI-like gene. At 5 hpi, only 2 genes, encoding a lebocin antibacterial (52) AMP (Lebocin 2) and the negative regulator Pirk of the Imd pathway (129), were found to be significantly upregulated. However, most of the selected genes that are strongly induced at later time points also presented positive log2 fold changes at this time point (Fig 2A). From 10 to 20 hpi, all selected genes but few exceptions (cecropin D, Tg-like and Integrin β-like) due to biological variability were significantly upregulated at each time point (Fig 2A). Clustering analyses based on Pearson coefficients however revealed 3 distinct clusters of covariations. The first one contained 13 genes belonging to all the categories cited above and corresponded to very stable induction patterns (Fig 2A). The second one, which contained 8 genes involved antimicrobial and cellular responses plus the selected C-type lectin (CLECT (ccBV)), corresponded to slightly increasing patterns (Fig 2A). Finally, the third one, which contained the Relish and Pelle members of the Imd and Toll pathways (22), an integrin and the DDC melanization enzyme (130) genes, corresponded to slightly decreasing patterns (Fig 2A).

**Figure 2.**
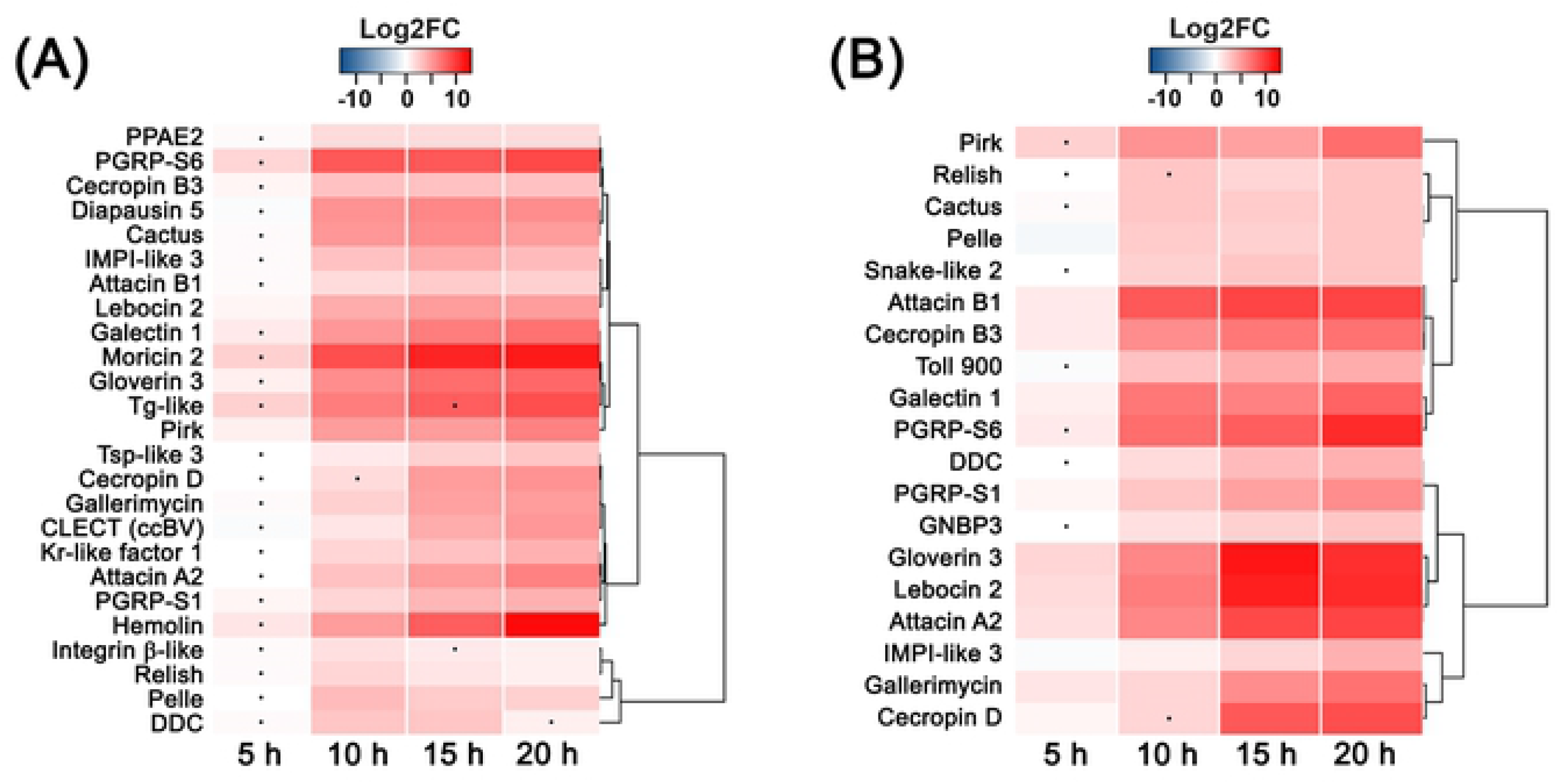
Temporal dynamics of the identified immune responses after infestation by the nematobacterial complex. Heatmaps showing the temporal evolution of the induction levels of representative immune genes in the hemocytes (A) and in the fat body (B) after infestation by the nematobacterial complex. RT-qPCR relative quantifications were performed on triplicate samples of N=9 larvae par sample with the RpL32 housekeeping gene as reference. Differential expression was assessed with Student t tests on ΔCq (149) and black dots indicate genes with statistically non-significant variations to the controls in the corresponding tissue (p-value > 0.05). The dendrograms represent clustering analyses based on Pearson correlation coefficients.

In the fat body, the selected genes included 15 genes of the antimicrobial response, 2 genes involved in melanization, one galectin gene (Galectin 1) and an IMPI-like gene (IMPI-like 3). At 5 hpi, all 7 selected AMPs, PGRP-S1 and Galectin 1 were found to be upregulated (Fig 2B). All these genes were among the most strongly overexpressed at later time points. Such as in the hemocytes, most of the selected genes were then significantly upregulated from 10 to 20 hpi (Fig 2B). In this tissue, the genes only subdivided into two main covariation clusters: a cluster of genes with stable induction patterns and a cluster of genes with increasing induction patterns. The first cluster contained 10 genes of which 8 were involved in antimicrobial responses, one encoded a melanization-related serine protease (Snake-like 2) and one encoded the Galectin 1 (Fig 2B). The second cluster contained 9 genes, of which 7 were involved in antimicrobial responses, one encoded the DDC melanization enzyme (130) and the last one encoded the IMPI-like 3 (Fig 2B).

Altogether, the results obtained for the two tissues show that most of the transcriptional immune responses induced at 15 hpi take place between 0 and 10 hpi, which is comparable to timings observed in other interaction models (131–133). The results also indicate that these responses are globally stable across the time post-infestation despite some distinct gene induction patterns in each category of response. Interestingly, while we were hoping to discriminate between an early response, probably activated by the nematode presence, and a later response, probably reacting to bacterial growth, we did not find any clear link between the gene inductions’ dynamics and the different immune processes and pathways that were represented in our selection.

### Evaluation of each NBC partner’s part in the induced immune responses

In order to identify each NBC partner’s relative participation in the fat body’s and hemocytes’ immune responses, we used RT-qPCR to compare the induction levels of the selected immune genes after independent infections by the whole NBC, the axenic nematode or the bacterial symbiont. To this aim, we decided to use a more standardized protocol of direct injection of the pathogens into the hemocoel, thereby limiting putative side effects such as early hemocoel colonization by intestinal microorganisms. Importantly, we previously compared the kinetics of *X. nematophila* growth and of *S. frugiperda* survival after injection of the entire NBC and of 200 *X. nematophila* (S2A and S2B Fig). This comparison showed that both kinetics are very similar and thus that any difference of induction level between the 2 conditions would not reflect differences in bacterial load or physiological state. However, the putative impact of axenization on the nematode’s physiology could not be assessed by the same way due to technical limitations and to its avirulence in absence of its bacterial symbiont (S2B and S2C Fig).

In the hemocytes, 14 genes presented higher induction levels in response to *X. nematophila* than in response to the axenic nematode (Fig 3). In the antimicrobial category, they included the negative regulator Pirk of the Imd pathway (129), all the selected attacin, cecropin, gloverin, lebocin and gallerimycin AMPs, the 2 selected PGRP-S, and also probably the Imd pathway transcription factor Relish (22) (Fig 3A). As indicated above, the Imd pathway, as well as the attacin, cecropin and gloverin AMP families, are known to take part in anti-Gram negative bacteria immune responses (11, 52). Their induction patterns thus indicate that the antimicrobial *X. nematophila*-induced responses are well adapted to the nature of the pathogen. Moreover, these results are in agreement with the study of Aymeric and colleagues (44) showing that the Imd pathway functions in the *D. melanogaster* immune response to *X. nematophila*. In the other categories, the *X. nematophila*-induced genes encoded the DDC melanization enzyme (130), the hemolin antibacterial opsonin (80), the IMPI-like 3, and also probably the selected integrin (Integrin β-like) (Fig 3B, 3C and 3D). Once again, all of these genes are susceptible to play a part in an immune response to a pathogenic bacterium even though most of them could act on diverse types of invaders. Surprisingly, we found that *X. nematophila* strongly over-induces the transglutaminase (Tg-like) putative clotting factor (85) (Fig 3C). This result could suggest that the bacterium is actually the main responsible for tissue damages at this time point and/or that Tg-like expression is induced in response to bacteria. Importantly, this result is in agreement with the study of Yadav and colleagues (43), who showed that the *D. melanogaster* Fondue clotting factor was induced after infestation by the NBC but not after infestation by axenic nematodes. Remarkably, most of the genes that were mostly induced by *X. nematophila* presented higher induction values in response to the bacterium alone than in response to the whole NBC. However, this observation cannot be directly interpreted as an antagonistic effect of the nematode partner since it could be due to changes in the relative proportions of each hemocyte subtype, which would not necessarily reflect absolute variations in their numbers. In addition, the nematode partner specifically induced the overexpression of the selected C-type lectin (CLECT (ccBV)) and was probably the main inducer of the Galectin 1, the tetraspanin D76 homolog (Tsp-like 3) and the selected diapausin AMP (Diapausin 5) (Fig 3A, 3C and 3D). As mentioned before, the *M. sexta* tetraspanin D76 is known to take part in encapsulation (83) and some lectins can bind nematodes and participate in melanization (109) as well as in all types of cellular immune responses. Once again, their induction patterns are consistent with the nature of the pathogen, since both types of molecules could be involved in classical anti-nematode immune responses, such as cellular or melanotic encapsulation (134). Finally, 5 genes, encoding the Toll pathway members Pelle and Cactus (22), the selected moricin AMP (Moricin 2), the melanization-related PPAE2 and the Krüppel-like transcription factor (Kr-like factor 1), were similarly induced by each of the three pathogens (Fig 3A, 3B and 3C), suggesting that these responses are induced by the 2 partners without any additive effect.

**Figure 3.**
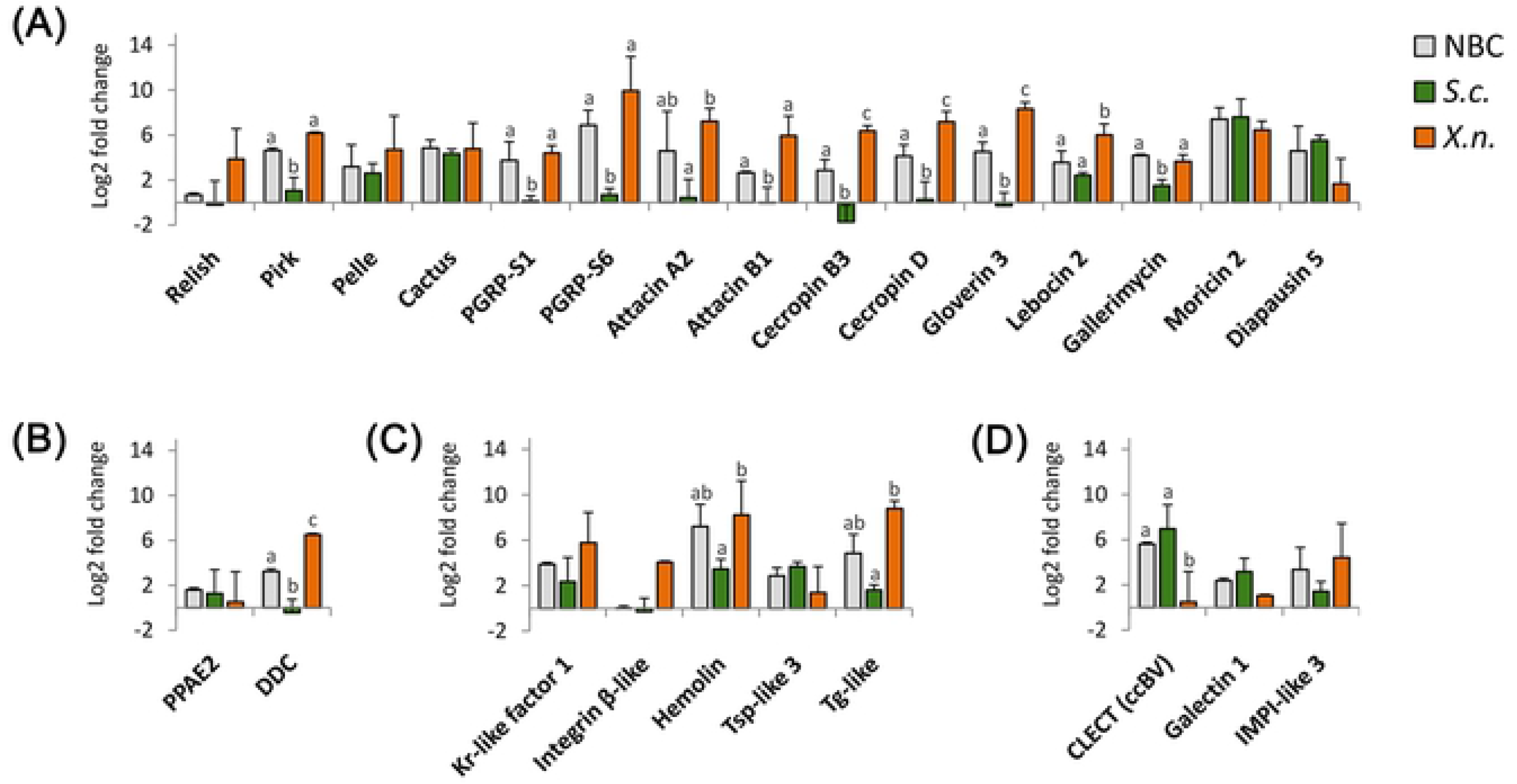
Relative participations of *S. carpocapsae* and *X. nematophila* in the hemocytes’ immune responses. Histograms showing the induction levels (+/− SEM) of representative immune genes in the hemocytes at 13 h after independent injections of either 10 nematobacterial complexes (NBC), 10 axenic *S. carpocapsae* (*S.c.*) or 200 *X. nematophila* symbionts (*X.n.*). RT-qPCR relative quantifications were performed on triplicate samples of N=9 larvae per sample with the RpL32 housekeeping gene as reference and buffer-injected control larvae. Letters show statistical differences between treatments from one-way ANOVA and Tukey tests on ΔΔCq (149). The genes were gathered by type of immune response with (A) antimicrobial immunity-related, (B) melanization-related, (C) cellular immunity-related and (D) diverse immune responses.

In the fat body, statistical analysis of the results firstly revealed that the induction levels of Pirk as well as of the selected cecropin and gloverin AMPs were significantly lower in response to the axenic nematode than in response to the NBC and to *X. nematophila* (Fig 4A), suggesting the bacterial partner is the main responsible for their inductions. In addition, despite non-significant statistics, the results for the selected attacin AMP, PGRP-S6 and GNBP3 showed similar induction patterns (Fig 4A). As for the hemocytes, the induction patterns of Pirk and of the attacin, cecropin and gloverin AMPs suggest that the fat body’s antimicrobial response to *X. nematophila* is well adapted to the type of pathogen that is met. On the contrary, the induction levels of the melanization-related serine protease (Snake-like 2) was significantly lower in response to *X. nematophila* than in response to the NBC and to the axenic nematode (Fig 4B), suggesting that the nematode partner is the main responsible for its induction. Similar induction patterns were obtained for the Toll pathway members Toll and Cactus (22) as well as for Galectin 1 (Fig 4A and 4C). As mentioned for the hemocytes, the induction of lectins and melanization-related genes in response to the nematode is consistent with the nature of the pathogen since both could participate in classical anti-nematode immune responses (134). The induction of Toll pathway members is more difficult to relate with known anti-nematode immune responses and Yadav and colleagues (47) found that the inactivation of this pathway does not impact the *D. melanogaster* survival to infestation by the whole NBC or by axenic *S. carpocapsae*. Therefore, the involvement of this immune pathway in anti-nematode immune responses may depend on the downstream effectors and thus be variable between insect species. Finally, the other genes did not show any clear difference of induction level after injection of the 3 pathogens, except for the gallerimycin AMP, PGRP-S1 and the DDC melanization enzyme, which presented a lesser induction when each NBC partner was injected alone (Fig 4A and 4B). These results suggest synergistic effects of the nematode and of the bacterium on the induction of these genes.

**Figure 4.**
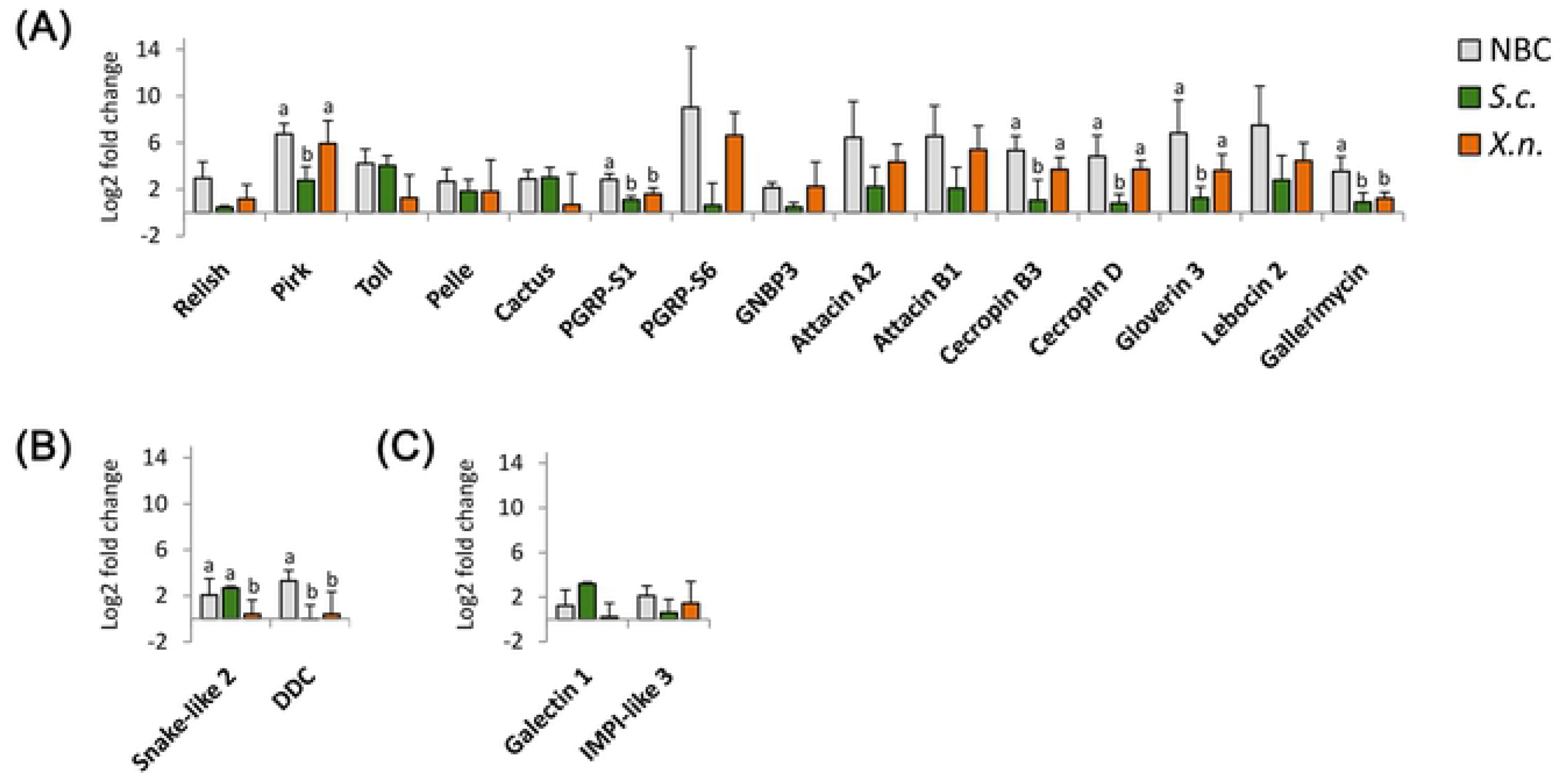
Relative participations of *S. carpocapsae* and *X. nematophila* in the fat body’s immune responses. Histograms showing the induction levels (+/− SEM) of representative immune genes in the fat body at 13 h after independent injections of either 10 nematobacterial complexes (NBC), 10 axenic *S. carpocapsae* (*S.c.*) or 200 *X. nematophila* symbionts (*X.n.*). RT-qPCR relative quantifications were performed on triplicate samples of N=9 larvae per sample with the RpL32 housekeeping gene as reference and buffer-injected control larvae. Letters show statistical differences between treatments from one-way ANOVA and Tukey tests on ΔΔCq (149). The genes were gathered by type of immune response with (A) antimicrobial immunity-related, (B) melanization-related and (C) diverse immune responses.

In summary, we found in the 2 tissues that most of the selected genes presented partner-specific induction patterns, suggesting that the immune response to the NBC corresponds to combinations of responses induced by each partner. The detailed analysis of these genes indicates that *X. nematophila* is the main inducer of most of the selected genes, and especially of the well-known antibacterial ones. On the other hand, *S. carpocapsae* is the main inducer of some melanization and encapsulation-related genes and of the selected lectins, which could all take part in classical anti-nematode immune responses. The results thus globally suggest that the hemocytes and the fat body both respond by adapted ways to each NBC partner despite some yet unexplained results, such as an induction of Toll pathway members in the fat body by the nematode partner.

### Expression patterns of two new clusters of candidate immune genes

During our first analysis of the RNAseq data, we identified 2 new clusters of candidate immune genes (48). The first one, named the Unknown (Unk) cluster, was localized close to Tamozhennic, a gene encoding a nuclear porin involved in the nucleation of Dorsal, the transcription factor of the Toll pathway (135). It contained 5 genes predicted to encode secreted peptides and short proteins that were all highly overexpressed in the midgut, fat body and hemocytes at 15 hpi and of which 4 were the unique mobilized genes at 8 hpi in the fat body. The second cluster, named the Genes with Bacterial Homology (GBH) cluster, contained 3 genes located inside a defensin-like AMP cluster in the *S. frugiperda* genome. The 3 genes were predicted to encode secreted proteins similar to each other and one of them was also found highly induced at 15 hpi in the 3 tissues. The particularity of these genes is that homologs are found only in lepidopteran species as well as, intriguingly, in Gram positive bacteria. Here, we reexamined the expression patterns of the Unk and GBH genes and found that the 5 Unk genes were mainly expressed in the fat body whereas 2 of the 3 GBH genes were mainly expressed and induced in the hemocytes (S2 Table). In order to learn more about their putative functions, we decided to analyse, as we did for the known immune genes, their induction patterns across the time post-infestation and in response to each NBC partner in the corresponding tissues. In both cases, we found that the induction dynamics of the genes were very similar to those of immune genes, with an upregulation that becomes significant at 5 or 10 hpi and with globally stable induction patterns from 10 to 20 hpi (Fig 5A and 5B).

**Figure 5.**
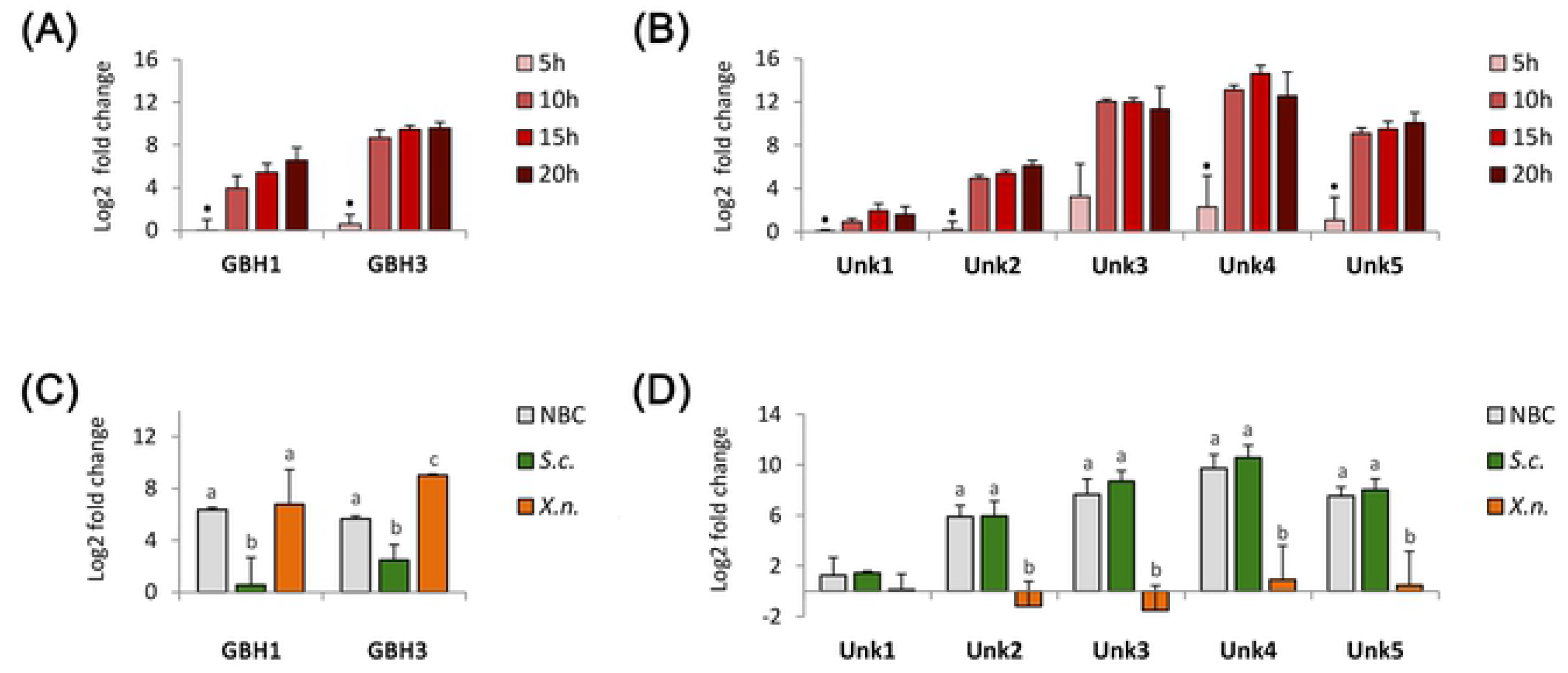
Transcriptional induction patterns of putative new immune genes. (A, C) Histograms showing the induction levels (+/− SEM) of 2 GBH genes in the hemocytes (A) and of the 5 Unk genes in the fat body (C) at several times after infestation by the nematobacterial complex. RT-qPCR relative quantifications were performed on triplicate samples of N=9 larvae per sample with the RpL32 housekeeping gene as reference. Differential expression was assessed with Student t tests on ΔCq (149) and black dots indicate genes with statistically non-significant variations to the controls (p-value > 0.05). (B, D) Histograms showing the induction levels (+/−SEM) of 2 GBO genes in the hemocytes (B) and of the 5 Unk genes in the fat body (D) at 13 h after independent injections either 10 nematobacterial complexes (NBC), 10 axenic *S. carpocapsae* (*S.c.*) or 200 *X. nematophila* (*X.n.*). RT-qPCR relative quantifications were performed on triplicate samples of N=9 larvae per sample with the RpL32 housekeeping gene as reference and buffer-injected control larvae. Letters show statistical differences between treatments from one-way ANOVA and Tukey tests on ΔΔCq (149).

In the case of the GBH cluster, the results that we got for the 2 NBC-responsive genes (GBH1 and GBH3) in the hemocytes indicate that they are significantly less induced after axenic nematode injection than after NBC and *X. nematophila* injections, suggesting that the bacterium is the main responsible for their up-regulation (Fig 5C). We could hypothesize an acquisition by horizontal gene transfer from bacteria of the GBH genes. In this case, their putative involvement in the antibacterial immune response would be particularly interesting, since bacterial genes hijacking for immune purpose has only been reported once in metazoans, in the tick *Ixodes scapularis* (136). Such a hypothesis however requires functional confirmation. In the case of the Unk cluster, we found that the 4 most induced genes in the fat body (Unk2 to 5) are all strongly and similarly induced by the NBC and by the axenic nematode whereas they are not induced by *X. nematophila* (Fig 5D). The results are very similar for the least expressed Unk gene (Unk1), for which we only found a significant induction for the injection of axenic nematodes (Fig 5D). This partner-specific induction pattern suggests the Unk genes are involved in specific aspects of the insect responses to the infestation. In addition, the putative involvement of the Unk genes in the response towards the nematode partner seems to be in agreement with their early mobilization during the infectious process and with their overexpression in the midgut, which is the entry site of the nematode. In our previous study, we had hypothesized the Unk may encode new types of immune effectors (48). However, given their low levels of conservation in species as close as *S. littura* or *S. littoralis* (S4 Fig) another hypothesis would be that they correspond to regulatory long non-coding RNAs (137, 138). In both cases, the further functional characterization of these genes could be very promising given our current lack of knowledge of the immune pathways and molecular effectors of insect anti-nematode immunity.

## Conclusion

Here, we provide a very deep and contextualized analysis of the *S. frugiperda*’s hemocytes’ and fat body’s transcriptional immune responses to infestation by the *S. carpocapsae-X. nematophila* NBC. Our topologic analysis of these responses at 15 hpi firstly confirmed the induction of very potent and diversified immune responses towards the pathogen, such as suggested by our previous analysis of the transcriptomic data (48) as well as by the study of Yadav and colleagues (43) in the *D. melanogaster* model. The present work establishes that these responses are very stable across the post-infestation time and that they correspond to combinations of *X. nematophila*- and *S. carpocapsae*-induced responses that seem to be well adapted to the nature of each partner (Fig 6).

**Figure 6.**
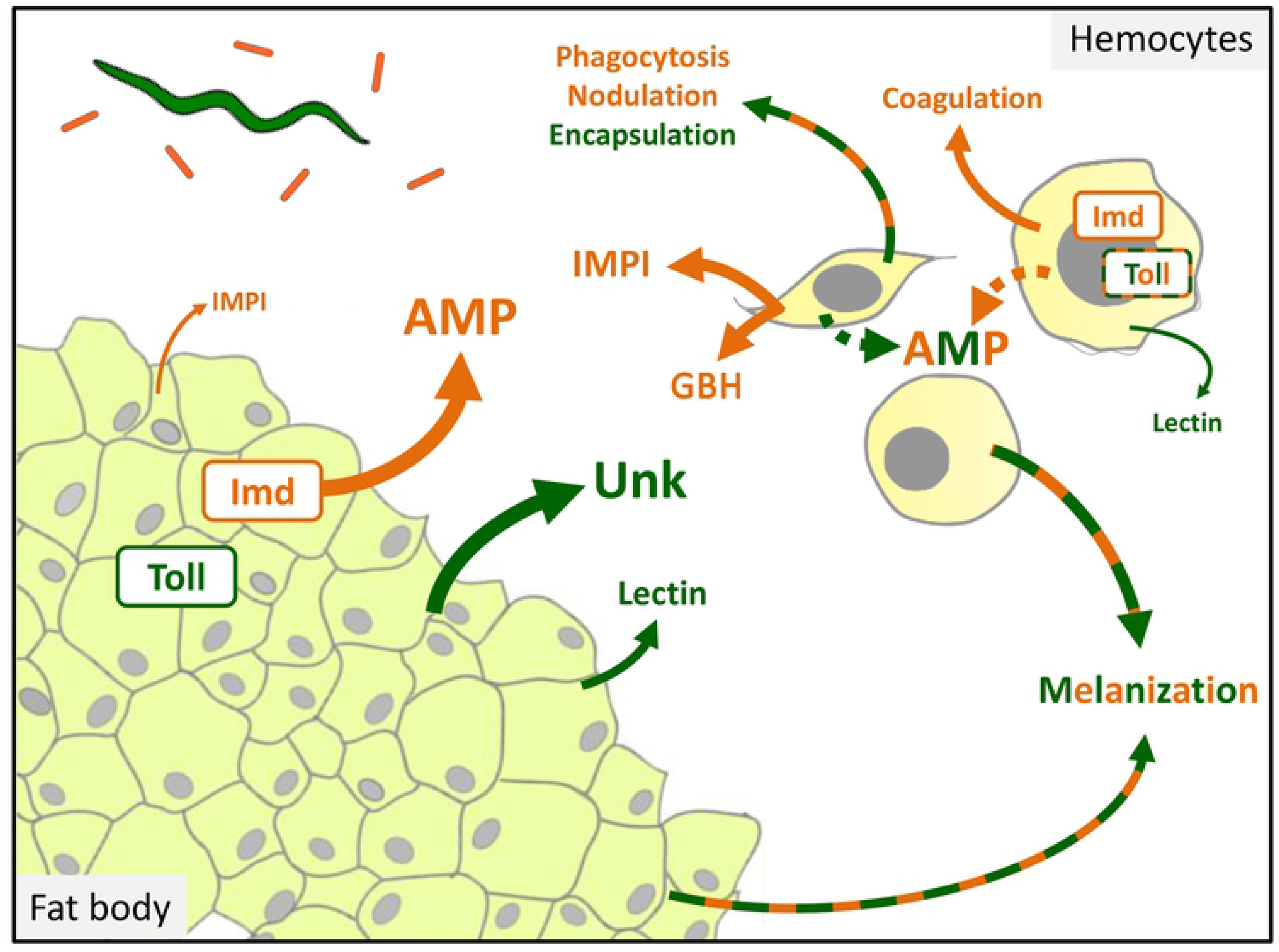
Hypothetical structure of the *S. frugiperda* larva’s immune response to the nematobacterial complex. Graphical abstract illustrating the main hypotheses we can emit from the present RNAseq and RT-qPCR data and from our current knowledge of *S. frugiperda* immunity. Dark green letters, lines and arrows indicate responses that seem to be mainly induced by the nematode partner *S. carpocapsae* whereas orange ones indicate responses that seem to be mainly induced by the bacterial symbiont *X. nematophila*. The arrows’ thicknesses and the letter sizes refer to the relative strengths of the induced transcriptional responses. AMP: AntiMicrobial Peptides, IMPI: Induced MetalloProteinase Inhibitors.

The pieces of information collected during these analyses are of great interest for the study of the dialogue that takes place between each NBC partner and their hosts’ immune systems. First, our results strongly suggest that the NBC immunosuppressive strategies globally have a low impact on the induction of immune responses at the transcriptional level. They also indicate that the nematode and/or its effects on the host are detected by the insect’s immune system that in return seems to induce adapted immune responses towards the pathogen. Such observations could help to identify the limits of previously described immunosuppressive and immunoevasive strategies of the NBC. For example, they suggest that the suppressive effect of *X. nematophila* on the expression of AMP genes (42, 60, 61) as well as the camouflage strategy of *S. carpocapsae* (24, 25) are probably far from sufficient to explain their success towards the immune system in the case of *S. frugiperda*. Nevertheless, we found several unexplained down-regulations of signaling genes, such as of members of the Imd, JNK, TGF-β and Jak-Stat pathways, that represent interesting working trails for the study of the molecular basis of the NBC’s immunosuppressive strategies. Finally, this study allowed the identification of very large panels of candidate immune genes involved in all the main components of insect immunity as well as of some yet uncharacterized genes that could encode new immune factors involved in the response to the complex.

Continuing this work with more functional and mechanistic approaches is now required to get an accurate picture of the molecular dialogue between the NBC and the immune system. In the longer term, such approaches could help to identify the precise causes of the immune system’s failure against this NBC and thus the conditions that are required for an adequate use of this NBC against insect pests.

## Materials and Methods

### Insect rearing

Corn variant *Spodoptera frugiperda* (Lepidoptera : Noctuidae) were fed on corn-based artificial diet (139). They were reared at 23°C +/− 1°C with a photoperiod of 16 h/8 h (light / dark) and a relative humidity of 40 % +/− 5 %. *Galleria mellonella* (Lepidoptera : Pyralidae) were reared on honey and pollen at 28°C in dark.

### Production and storage of nematobacterial complexes

*Steinernema carpocapsae-Xenorhabdus nematophila* complexes (strain SK27 isolated from Plougastel, France) were renewed by infestation of one month-old *Galleria mellonella* larvae. They were collected on White traps (140) and stored at 8°C in aerated Ringer sterile solution with 0.1 % formaldehyde. The maximal time of storage was limited to 4 weeks to avoid pathogenicity losses.

### Production of axenic nematodes

Gravid *S. carpocapsae* females were extracted from *G. mellonella* dead bodies at day 4 to 6 after infestation by nematobacterial complexes. After 5 washing steps in Ringer sterile solution, the females were surface-sterilized by 20 min incubation in 0.48% (wt/vol) sodium hypochlorite and 3 h incubation in Ringer sterile solution supplemented with antibiotics (150 μg/mL polymyxin, 50 μg/mL colistin, 50 μg/mL nalidixic acid). The eggs were extracted by female crushing with sterile glass pestles and then washed by centrifugation (2 min, 16000 g) in Ringer sterile solution, disinfected by incubation in 0.48% sodium hypochlorite for 5 min, and washed again twice. After microcopic observation, the intact eggs were placed on liver-agar (40 g/L Tryptycase Soja Agar [BioMérieux], 5 g/L Yeast Extract [Difco], 100 g/L porc liver) plates supplemented with antibiotics (150 μg/mL polymyxin, 50 μg/mL colistin and 50 μg/mL nalidixic acid). The plates were maintained inside a dark humid chamber for 1 month to allow nematodes development. The nematodes were then suspended in Ringer sterile solution and infective juvenile stages (IJs) were sorted by pipetting under a microscope (Leica). The IJs were rinsed twice by centrifugation (2 min, 3000 g) in 1 mL Ringer sterile solution and used within minutes for experimental infection.

Nematodes’ axenicity was verified *a posteriori* by DNA extraction and PCR amplification. Nematodes were suspended in 200 μL milliQ water supplemented with 200 μL glass beads (Ø ≤ 106 μm) (Sigma). They were grinded for 2 × 40 sec at 4.5 ms speed with a FastPrep homogenizer (MP Biomedicals). The debris were discarded by centrifugation (2 min, 16000 g) and 150 μL supernatant were mixed with 200 μL lysis buffer (Quick extract kit, Epi-centre) for a second grinding. To ensure bacterial cell lysis, the samples were incubated at room temperature for 48 h with 2 μL Ready-Lyse Lysozyme solution at 30000 U/μL (Epi-centre). Protein denaturation was then performed by 10 min incubation at 90°C, and RNA was removed by 10 min incubation at 37°C with 20 μL RNase A (20 mg/mL) (Invitrogen). DNA was extracted by successive addition of 500 μL phenol-chloroform-isoamyl alcohol and 500 μL chloroform, followed by centrifugations (10 min, 16000 g) and aqueous phase collections. DNA was precipitated with 500 μL 100% ethanol supplemented with 20 μL sodium acetate and by freezing at −80°C for 2 h. After defrosting, DNA was concentrated by centrifugation (30 min, 16000 g) and the precipitates were washed twice by centrifugation (15 min, 16000 g) in 500 μL 70% ethanol. DNA was finally suspended in 50 μL sterile milliQ water and left at room temperature for a few hours to ensure precipitate dissolution. After DNA quantification with a Qubit fluorometer (Invitrogen), *X. nematophila* presence was assessed by PCR amplification with *Xenorhabdus*-specific primers (Xeno_F: 5’-ATG GCG CCA ATA ACC GCA ACT A-3’; Xeno_R: 5’-TGG TTT CCA CTT TGG TAT TGA TGC C-3’), which target a region of the XNC1_0073 gene encoding a putative TonB-dependent heme-receptor. The presence of other bacteria was assessed by 16S rRNA gene amplification with universal primers (141). Thirty cycles of PCR were performed using Taq polymerase (Invitrogen) in a Biorad thermocycler (Biorad), with hybridization temperatures of 55°C and 50°C respectively. PCR products were then analyzed by agarose gel electrophoresis.

### Experimental infections

Experimental infestations with nematobacterial complex were carried out on individual 2nd day 6th instar *S. frugiperda* larvae according to (48). Larvae were kept at 23°C in 12-well plates with an articial diet (139). Briefly, each well was coated with a piece of filter paper (Whatman) and 150 +/− 20 NBCs in 150 μL Ringer solution were poured in each larva-containing well. 150 μL Ringer sterile solution were used for control larvae.

For intra-hemocoelic injection experiments, pathogens were injected in larvae’s abdomens after local application of 70% ethanol with a paintbrush. Injections were performed using a syringe pump (Delta labo) with 1 mL syringes (Terumo) and 25G needles (Terumo). *X. nematophila* suspensions were prepared as described in Sicard et al (2004)(142). Bacterial culture was diluted in PBS and 20 μL containing 200 +/− 50 bacterial cells were injected in the hemocoel at a rate of 1.67 mL/min. 20 μL sterile PBS was used for control larvae. The purity and number of injected *X. nematophila* were verified by plating 20 μL of the bacterial suspension on NBTA (143). For NBC and axenic nematode injections, 10 +/− 3 nematodes in 20 μL solution at 70% Ringer and 30% glycerol were injected at a rate of 2.23 mL/min. Syringes were frequently renewed in order to limit nematodes’ concentration and sedimentation and the number of injected nematodes was verified by 10 simulations of injection in Petri dishes followed by nematode counting under a microscope (Zeiss). Sterile solutions at 70% Ringer and 30% glycerol were used for control larvae. To avoid accidental *per os* infections, the injected larvae were then briefly washed in sterile PBS and dried on paper towel before being placed in 12-well plates. The pathogens efficacies were checked by monitoring 12 control and 12 infected larvae’s survival for 72 h after infestation or after injection.

### Production and storage of bacterial symbionts

*X. nematophila* strain F1 isolated from nematobacterial complexes strain SK27 was conserved at −80°C. Within 3 weeks before each experiment, they were grown for 48 h at 28°C on NBTA with erythromycin (15 μg/mL). The colonies were then conserved at 15°C and used for overnight culture at 28°C in 5 mL Luria-Bertani broth (LB) before experiments.

### RNA extraction

RNAs were prepared as described in Huot et al (2019) (48). Briefly, nine larvae per technical replicate were bled in anti-coagulant buffer (144). Hemocytes were recovered by centrifugation (1 min, 800 g) at 4°C and the pellet was immediately flash-frozen with liquid nitrogen. The larvae were then dissected for fat body and midgut sampling and the tissues were flash-frozen in eppendorf tubes with liquid nitrogen. After storage at −80°C for at least 24 h, 1 mL Trizol (Life technologies) was added to the pooled tissues. The tissues were then grounded by using a TissueLyzer 85210 Rotator (Qiagen) with one stainless steel bead (Ø : 3 mm) at 30 Hz for 3 min. For optimal cell lyses, grounded tissues were left at room temperature for 5 min. To extract nucleic acids, 200 μL chloroform (Interchim) were added and the preparations were left at room temperature for 2 min with frequent vortex homogenization. After centrifugation (15 min, 15,000 g) at 4°C, the aqueous phases were transferred in new tubes and 400 μL 70% ethanol were added. RNA purifications were immediately performed with the RNeasy mini kit (Qiagen) and contaminant DNA was removed with the Turbo DNA-freeTM kit (Life Technologies).

RNA yield and preparation purity were analyzed by measuring the ratios A_260_/A_280_ and A_260_/A_230_ with a Nanodrop 2000 spectrophotometer (Thermo Scientific). RNA integrity was verified by agarose gel electrophoresis and RNA preparations were conserved at −80 °C.

### RNAseq experiments

RNAseq raw data originate from Huot et al (2019) (48). In brief, libraries were prepared by MGX GenomiX (IGF, Montpellier, France) with the TruSeq Stranded mRNA Sample preparation kit (Illumina). The libraries were then validated on Fragment Analyzer with a Standard Sensitivity NGS kit (Advanced Analytical Technologies, Inc) and quantified by qPCR with a Light Cycler 480 thermal cycler (Roche Molecular diagnostics). cDNAs were then multiplexed by 6 and sequenced on 50 base pairs in a HiSeq 2500 system (Illumina) with a single-end protocol. Image analysis and base calling were performed with the HiSeq Control and the RTA softwares (Illumina). After demultiplexing, the sequences quality and the absence of contaminant were checked with the FastQC and the FastQ Screen softwares. Data were then submitted to a Purity Filter (Illumina) to remove overlapping clusters.

For each sample, the reads were pseudoaligned on the *S. frugiperda* reference transcriptome version OGS2.2 (49) using the Kallisto software (145). Differential expression between infested and control conditions were then assessed for each time point and tissue with the Sleuth software (146). Wald tests were used with a q-value (equivalent of the adjusted p-value) threshold of 0.01 and a beta value (biased equivalent of the log2 fold change) threshold of 1. Only transcripts with normalized counts over 5 in all three replicates of the infested and/or of the control condition were considered as reliably differentially expressed.

Previously annotated immune transcripts (49) were then checked for significant expression changes and not annotated differentially expressed ones were researched with the Blast2GO software by blastx on the NCBI nr and drosophila databases (147). To avoid mistakes related to genome fragmentation, the immune transcripts were gathered by unique gene after careful examination of their sequences and of the available genomic data (49). The induction levels of the transcripts were then averaged by unique gene before graphical representation of the results.

### RT-qPCR experiments

cDNAs were synthesized from 1 μg of RNA with the SuperScript II Reverse Transcriptase (Invitrogen), according to the manufacturer’s protocol.

The primers (S3 Table) were designed with the Primer3Web tool (148). Their efficiency was estimated by using serial dilutions of pooled cDNA samples and their specificity was verified with melting curves analyses. Amplification and melting curves were analyzed with the LightCycler 480 software (Roche Molecular diagnostics).

RT-qPCR were carried out in triplicate for each biological sample, with the LightCycler 480 SYBR Green I Master kit (Roche). For each sample and primer pair, 1.25 μL of sample containing 50 ng/μL of cDNA and 1.75 μL of Master mix containing 0.85 μM of primers were distributed in multiwell plates by an Echo 525 liquid handler (Labcyte). The amplification reactions were then performed in a LightCycler 480 thermal cycler (Roche) with an enzyme activation step of 15 min at 95°C, and 45 cycles of denaturation at 95°C for 5 sec, hybridization at 60°C for 10 sec and elongation at 72°C for 15 sec.

Crossing points were determined using the Second Derivative Maximum method with the LightCycler 480 software (Roche) and relative expression ratios between control and infected conditions were manually calculated according to the method of Ganger et al (2017)(149). The ratios were normalized to RpL32 housekeeping gene relative levels and the EF1 gene was used as an internal control.

Statistical analyses of the data were all performed with the R software (150). Differential expression significance between the control and infected conditions was assessed by paired one-tailed t-tests on ΔCq values. Multiple comparisons of fold changes were assessed by one-way ANOVA on ΔΔCq values followed by post hoc Tukey tests. P-values under 0.05 were considered as significant for all the above tests. The gplots package was used to draw the heatmaps and the clusters were built from a dissimilarity matrix based on Pearson correlation coefficients.

### Quantification of nematodes in the midgut lumen

NBCs in the midgut lumen were quantified at several times after infestation by nematode counting in the alimentary bolus. For 3 independent experiments, 3 infested larvae were dissected and the midguts alimentary bolus were extracted. Each alimentary bolus was then dissolved in 3 mL sterile PBS in a Petri dish (Ø : 35 mm) and motile nematodes were counted with a microscope (Leica).

### Quantification of *X. nematophila* in the hemolymph

The concentration of *X. nematophila* in the hemolymph was estimated by CFU counting. For 3 independent infection experiments and 3 technical replicates, hemolymph was collected by bleeding of 3 caterpillars in 200 μL PBS supplemented with phenylthiourea (Sigma). The volumes of hemolymph were then estimated by pipetting and serial dilutions of the samples were plated on NBTA with 15 μg/mL erythromycin. CFU were counted after 48 h incubation at 28°C and the counts were reported to the estimated hemolymph volumes in order to calculate the bacterial concentrations. Hemolymph of naive larvae was also plated for control.

### Insect survival kinetics

Survival kinetics were performed in triplicate on pools of 20 infested or injected larvae. Survival was monitored from 0 to 72 hours after contact or injection. Naïve larvae were used as control for infestations whereas larvae injected with PBS were used for controls of *X. nematophila* injections and larvae injected with 70% Ringer − 30% glycerol solutions were used for controls of nematobacterial complexes and nematodes injections.

### Parasitic success measurement

Parasitic success was measured in triplicate on pools of 20 nematobacterial complexes or axenic nematodes-injected larvae. Dead larvae were individually placed on white traps (140) approximately 2 days after their deaths. The emergence of nematodes was assessed at day 40 after injection by observation of the collection liquid with a microscope (Leica). Parasitic success was then calculated as the percentage of larvae with nematode emergence among the infected larvae.

## Acknowledgments

We thank the quarantine insect platform (PIQ), member of the Vectopole Sud network, for providing the infrastructure needed for pest insect experimentations. We are also grateful to Clotilde Gibard and Gaëtan Clabots for maintaining the insect collections of the DGIMI laboratory in Montpellier. This work was supported by grants from the French Institut National de la Recherche Agronomique.

## Authors’ contribution

L.H., N.N. and B.D. conceived this study. N.N. and B.D. directed this study. L.H. and P.-A.G. performed the infestation experiments. L.H., P.-A.G. performed dissections. L.H. and A.B. extracted and purified the RNA. J.-C.O. designed the *X. nematophila* specific primers. S.P. produced the axenic nematodes and checked their axenization. L.H. and A.B. performed the qPCRs. L.H., N.N. and B.D. analysed the data. L.H. wrote the manuscript. L.H., N.N. and B.D. revised the manuscript. All authors have read and approved the manuscript.

## Supporting Information Legends

**S1 Table. Hemocytes and fat body RNAseq results for *S. frugiperda*’s immune genes.** (A) Results for the previously annotated *S. frugiperda*’s immune genes, (B) Results for the newly identified *S. frugiperda*’s immune genes. The statistics of the transcripts that were considered as significantly (Sleuth : |Beta|>1; qval < 0.01; pseudocounts > 5 in all the samples of at least one condition) up- or down-regulated are highlighted in red and blue, respectively. The Beta value gives a biased estimate of the log2 fold change. The qvalue (qval) is an equivalent of the adjusted p-value. The following columns give the normalized pseudocounts (Kallisto) for each individual sample, with HCn15 and FBn15 corresponding to control larvae and HCi15 and FBi15 corresponding to infested larvae. Blast hits on the *Drosophila* and nr NCBI databases were obtained by blastx with the Blast2GO sofware.

**S2 Table. Hemocytes and fat body RNAseq results for the Unk and GBH putative new immune genes.** The statistics of the transcripts that were considered as significantly upregulated (Sleuth : Beta>1; qval < 0.01; pseudocounts > 5 in all the samples of at least one condition) are highlighted in red. The Beta value gives a biased estimate of the log2 fold change. The qvalue (qval) is an equivalent of the adjusted p-value. The following columns give the normalized pseudocounts (Kallisto) for each individual sample, with HCn15 and FBn15 corresponding to control larvae and HCi15 and FBi15 corresponding to infested larvae.

**S3 Table. Primers sequences and genes used in this study.**

**S1 Fig. Temporal monitoring of nematobacterial infestation parameters.** (A) Dotplot showing the number of *S. carpocapsae* detected in the midgut alimentary bolus at several times after infestation by the nematobacterial complex. Infestations were performed by putting in contact individual larvae with 150 nematobacterial complexes (at time 0) in cell culture plates. Dot colors correspond to 3 independent experiments on N=3 larvae per time point. (B) Curve showing the temporal evolution of *X. nematophila* concentration (+/−SEM) in the hemolymph across the time post-infestation. Infestation experiments were performed in triplicate with 3 pools of 3 larvae per time point. *X. nematophila* were quantified by CFU counting on selective culture medium. (C) Curve showing the temporal evolution of *S. frugiperda* larvae’s survival percentage (+/− SEM) across the time post-infestation. Infestation experiments were performed in triplicate on N=20 larvae per experiment.

**S2 Fig. Comparison of the main infection parameters after independent injections of the nematobacterial complex, of axenic *S. carpocapsae* and of *X. nematophila*.** (A) Curves showing the temporal evolution of *X. nematophila* concentration (+/−SEM) after independent injections of either 10 nematobacterial complexes (NBC) or 200 *X. nematophila* (*X.n.*). Injection experiments were performed in triplicate with 3 pools of 3 larvae per time point and *X. nematophila* were quantified by CFU counting on selective culture medium. (B) Curves showing the temporal evolution of S*. frugiperda* larvae’s survival percentage (+/− SEM) after independent injections of either 10 nematobacterial complexes (NBC), 10 axenic *S. carpocapsae* (*S.c.*) or 200 *X. nematophila* (*X.n.*). Injection experiments were performed in triplicate on N=20 larvae per experiment. No insect death was reported for control buffer-injected larvae. (C) Histogram showing the parasitic success (+/− SEM) (i.e.: number of larvae with nematobacterial complex emergence on total number of infested larvae) after independent injections of either 10 nematobacterial complexes (NBC) or 10 axenic *S. carpocapsae* (*S.c.*). Injection experiments were performed in triplicate on N=20 larvae per experiment.

**S3 Fig. Verification of *S. carpocapsae* axenicity.** Electrophoresis gel showing the absence of bacterial contaminants in the axenized nematodes. Total DNAs from grinded infective stage nematodes (axenic *S.c.)* were extracted and the absence of bacterial contaminants was verified by PCR amplification of the 16S rRNA gene with universal primers and of the *Xenorhabdus*-specific gene (see Materials and Methods). Whole nematobacterial complexes (NBC) and a pure suspension of *X. nematophila* (*X.n.*) were used as positive controls. A pure suspension of *P. protegens* (*P.p.*) was used as negative control for putative TonB-dependent heme-receptor amplification.

**S4 Fig. Alignment of deduced amino acid sequences of Unks from *S. frugiperda* with those of *S. litura* and *S. littoralis*.** Nucleotide sequences were retrieved by blastn on *S. litura* and *S. littoralis* genomes.

